# IRE1-dependent GOLIM4 expression controls protein secretion to modulate glioblastoma cell adhesion and migration

**DOI:** 10.1101/2024.10.22.619629

**Authors:** Ketsia Bakambamba, Manon Nivet, Chloé Sauzay, Sophie Martin, Elodie Lafont, Luc Négroni, Eric Chevet, Tony Avril

**Affiliations:** INSERM UMR1242, Rennes, France; Centre Eugène Marquis, Rennes; IGBMC CNRS UMR7104, Illkirch, France

**Keywords:** glioblastoma, ER stress, IRE1, GOLIM4, GPP130, GOLPH4, protein secretion, cell adhesion, cell migration

## Abstract

One of the main glioblastoma (GB) features is the diffuse migration of the tumor cells within the surrounding brain parenchyma, rendering almost impossible the complete tumor resection and irradiation, leading to inexorable lethal relapse of the disease. In the past years, we demonstrated that IRE1α (hereafter IRE1), one of the Endoplasmic Reticulum (ER) stress sensors, plays a key role in GB biology by impacting on immune infiltration, angiogenesis and tumor cell migration/invasion, all these features being linked to an alteration of protein secretion. In the present study, we investigated if and how IRE1 could regulate the functionality of the secretory machinery in GB cells and identified GOLIM4, a Golgi-associated molecule whose expression is regulated downstream of IRE1 through the transcription of XBP1s. Interestingly, GOLIM4 silencing led to decreased surface expression of multiple molecules including MHC class I molecules, growth factor receptors (PDGFRA and IL13RA2) and proteins involved in cell-cell adhesion (CD44, CD54, NCAM1), adhesion to matrix (ITGB1) or cell migration (CD90) without alteration of their encoding transcripts’ expression levels. Moreover, GOLIM4 silencing phenotypically affected GB cell-cell adhesion and cell migration in multiple models. Overall, we have described a novel IRE1/XBP1s/GOLIM4 operon that controls the secretion of specific proteins and impacts the tumor aggressiveness.

**Graphical abstract:** 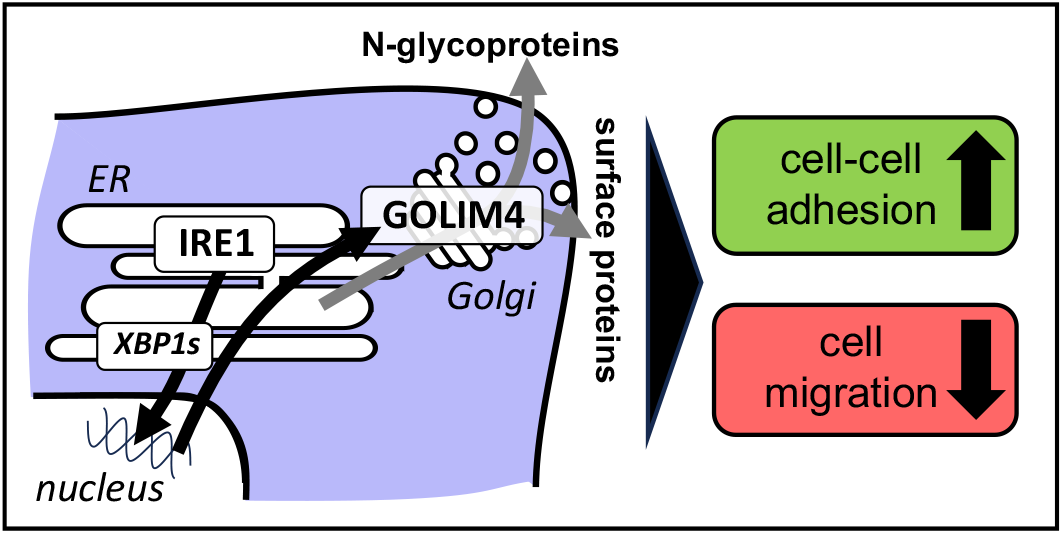

## Introduction

Upon oncogenesis, cells are exposed to acute demands of protein synthesis, that occur despite an unfavorable cell environment such as oxygen or nutrient deprivation. This defect in protein production in the Endoplasmic Reticulum (ER) could lead to an accumulation of unfolded or misfolded proteins inducing an ER stress (1,2). To restore ER homeostasis, ER stress induced cells display a signaling program leading to a transcriptional response called the unfolded protein response (UPR) (1–3). In mammals, three ER-resident proteins act as ER stress sensors ATF6, IRE1α (referred to as IRE1 hereafter) and PERK to initiate the UPR (1,2). One of these ER stress sensors IRE1 plays a key role in glioblastoma (GB) biology (4), affecting multiple cellular functions including immune recruitment (5,6), angiogenesis (7,8) or tumor cell migration (6,9).

The ER stress response, which triggers the activation of the Unfolded Protein Response (UPR) with IRE1 as one of its main transducers, aims at restoring ER homeostasis when perturbed thus ensuring proper function of the secretory pathway (SP). The SP is essential to maintain cellular functions and multicellular communications. The SP is executed through the intervention of several cell organelles (ER, Golgi apparatus, vesicles, …) and multiple molecular machineries (ER translocation, ER quality control and folding, ERAD, export, …) (10), all of those being challenged in cancer cells (11). Previous studies describe modifications of SP-associated organelles morphology upon ER stress, i.e. enlargement of ER cisternae (12,13), generation of new ER sheets and tubules via ER expansion (14–17), ER Golgi intermediate compartment (ERGIC) and Golgi apparatus fragmentation (18,19). The activation of the ER stress sensors also control the expression of molecular actors of the SP. For instance, upon prolonged ER stress, ER forms multimembrane vesicles called ER whorls initiated by SAR1 and SEC22 molecules, involving PERK activity (20). The cargo receptor ERGIC53 that recycles between the ER and the ERGIC to sort ER resident proteins is regulated by ATF6 (21). Moreover, IRE1, through the induction of the transcription factor XBP1s, induces the expression of the COPII components SEC22, SEC23 and SEC24 which are involved in the export of secreted proteins from the ER to the Golgi (22,23). Cargo receptors such as KDELR are also induced by the activation of the IRE1/XBP1s branch (23,24). Furthermore molecules of the COPI machinery i.e. COPA, COPB1/2, COPE and COPG involved in the retrograde protein trafficking are induced after XBP1s activation (23).

Recently, we have shown that IRE1-mediated via UBE2D3 expression control leads to NFκB activation and triggers proinflammatory chemokine secretion and the subsequent recruitment of immune/inflammatory cells to the tumor site in GB (5). In the present study, we aimed at investigating if and how IRE1 could more globally regulate the functionality of the secretory machinery in GB cells.

## Materials and Methods

### Antibodies and other reagents

Primary antibodies are listed in **Table S1**. Secondary antibodies used were horseradish peroxidase conjugated polyclonal goat anti-rabbit IgG, goat anti-mouse IgG and rabbit anti-goat IgG (all from Dako). Unless specified, all other reagents were from Sigma Aldrich.

### Cell culture and treatments

The immortalized GB cell U251 and U87 lines (all from ATCC) were grown in DMEM medium (Invitrogen) supplemented with 10% fetal bovine serum (FBS) (Lonza) in a 5% CO_2_ humidified atmosphere at 37°C. Primary GB lines RADH85 and RADH87 were previously described (25,26). For transient silencing of GOLIM4, IRE1 and XBP1, cells were transfected with specific siRNA (from Ambio, Santa Cruz and Dharmacon respectively) using Lipofectamine RNAiMAX Transfection Reagent (ThermoFisher Scientific), according to the manufacturers’ instructions as described in (5). Forty-eight to seventy-two hours after transfection, cells were used in either flow cytometry, Q-PCR, western blot or functional experiments (proliferation, adhesion or migration assays). For treatment with the IRE1 inhibitor MKC8866 (Selleckchem, Cologne, Germany), cells were incubated in culture medium in the presence of MKC8866 at 30 µM for 72 hours.

### Gene expression data analysis

Transcriptomes of parental and IRE1-modulated GB cell lines were previously described in (5,6). For tumor transcriptome derived from GB patients, the TCGA cohort was analyzed (https://www.cancer.gov/tcga). Complete gene expression analysis of the GB TCGA was performed with R (R version 3.5.0) / Bioconductor software as described in (5,6). Hierarchical clustering of GB patients labeled for their IRE1 activity (high or low), based on their ranked IRE1sign38 score obtained from (6), was coupled with GB patients survival and the gene of interest expression.

### Quantitative real-time PCR

Total RNA from GB cells was extracted using the TRIzol reagent (Invitrogen). RNAs were reverse-transcribed with Maxima Reverse Transcriptase (ThermoFisher) according to the manufacturer’s protocol. qPCR was performed with a QuantStudio™ 5 Real-Time PCR System and the PowerUp™ SYBR Green Master Mix (ThermoFisher). Experiments were performed at least in triplicate for each data point. Each sample was normalized based on the expression of the GAPDH or ACTB gene using the 2^ΔΔCT^-method. Primers pairs used in this study are listed in **Table S2**.

### Mass spectrometry

Parental U251, RADH85, RADH87 treated or not with the IRE1 inhibitors MKC8866 or Z4 (27), and IRE1 dominant negative U251_DN, RADH85_Q* and RADH87_Q* cells were lysed with lysis buffer composed of 20 mM Tris pH 8, 1.5 mM EDTA, 150 mM NaCl, 1% Triton X-100, 0.1% SDS, 15µM MG132, 10mM NEM (N-ethylmaleimide), supplemented with proteases and phosphatases inhibitor cocktails (Roche). Total proteins were precipitated overnight with 80% ice-cold acetone. Protein pellets were then washed 3 times with 80% acetone, followed by centrifugation at 500 g for 30 mins at 4°C. Samples were alkylated and digested with trypsin at 37°C overnight. After Sep Pak desalting, peptides were analyzed using an Ultimate 3000 nano-RSLC (Thermo Fisher Scientific) coupled in line with an Orbitrap ELITE (Thermo Scientific). Each sample was analyzed in triplicate. Briefly, peptides were separated on a C18 nano-column with a linear gradient of acetonitrile and analyzed in a Top 20 CID (Collision-induced dissociation) data-dependent mass spectrometry. Data were processed by database searching against Human Uniprot Proteome database using Proteome Discoverer 2.2 software (Thermo Fisher Scientific). Precursor and fragment mass tolerance were set at 7 ppm and 0.6 Da respectively. Trypsin was set as enzyme, and up to 2 missed cleavages were allowed.

Oxidation (M, +15.995), GG (K, +114.043) were set as variable modification and Carbamidomethylation (C) as fixed modification. Proteins were filtered with False Discovery Rate <1% (high confidence). Lastly, quantitative values were obtained from Extracted Ion Chromatogram (XIC) and p-values were determined by ANOVA with Precursor Ions Quantifier node in Proteome Discoverer.

### Western blotting

Cells were lysed in ice-cold lysis buffer (30 mmol/L Tris-HCl, pH 7.5, 150 mmol/L NaCl, 0.5% Triton X). Proteins were resolved by 10% SDS-PAGE and transferred to nitrocellulose membrane for blotting. The membranes were blocked with 5% BSA in 0.1% Tween 20 in PBS and incubated with the diluted primary antibodies (1/1000) (**Table S1**). Antibody binding was detected with the appropriate horseradish peroxidase-conjugated secondary antibodies (1/7000) and visualized with ECL (KPL, Eurobio) according to the manufacturer’s instructions and chemiluminescence signal was detected using G:Box Chemi XX6 imager from Syngene. Protein expression levels were determined by analyzing the luminescence signals using Fiji (28).

### Flow cytometry

Cells were washed in PBS 2% FBS and incubated with saturating concentrations of human immunoglobulins and fluorescent-labelled primary antibodies (**Table S1**) for 30 minutes at 4°C. Cells were then washed with PBS 2% FBS and analyzed using the Novocyte 3000 flow cytometer (Acea Biosciences).

### Cell-cell adhesion

Cell-cell adhesion was tested in cell aggregation experiments. Parental, controls, and U251 and U87 cells silenced for GOLIM4 (5,000 cells) were incubated in a 25 µL-drop on a cover of a 24-well plate. Images of cell aggregates were taken after 72 hours and cell density was estimated using Fiji by calculating the aggregates’ size. High aggregate’s size corresponds to the low cell-to-cell adhesion.

### Boyden chamber migration assay

Parental, controls, and U251 and U87 cell lines silenced for GOLIM4 were washed in DMEM, placed in Boyden chambers (10^5^ cells/chamber in DMEM) that were placed in DMEM 20% FBS and incubated at 37°C for 24 hours. After 24 hours, Boyden chambers were washed in PBS and cells were fixed in PBS 0.5% paraformaldehyde. Cells that did not migrate and remained inside the chambers were removed and cells that migrated were then stained with Giemsa (RAL Diagnostics). After washes in PBS, pictures of five different fields were taken. Migration index was given by the mean of number of migrated cells observed per field. ***Statistical analyses –*** Graphs and statistical analyses were performed using GraphPad Prism 7.0 software (GraphPad Software). Data are presented as the mean ± SD or SEM of at least three biological replicates. Statistical significance was determined using a paired, unpaired t-test or ANOVA as appropriate. Significant variations are represented by asterisks above the corresponding bar when comparing test and control conditions, and above the line when comparing the two indicated conditions.

Additional information can be found in the Supplemental Materials and Methods.

## Results

### IRE1 endoribonuclease activity modulates the secretion of N-glycosylated and cell surface proteins in GB cells

We recently demonstrated that IRE1 activity controls GB biology by secreting chemokines leading to the recruitment of inflammatory cells such as myeloid cells (5). Herein, we tested whether this was also true for other secreted proteins that could affect other GB cell functions. To this end, we investigated whether IRE1 modulation impacted on protein secretion i.e. N-glycoproteins (**Fig. 1A**) and surface proteins (**Fig. 1B**) using two different proteomic approaches. We first compared global N-glycoproteome of parental and GB cells in which IRE1 was inhibited using a dominant negative form of IRE1 or the IRE1 inhibitors MKC8866 and Z4 (**Fig. 1A**) we have previously described (5,6,27). Genetic ablation of IRE1 in RADH85/87 and U251 cells led to expression attenuation of 208 N-glycoproteins mainly involved in cell adhesion and migration whereas treatment with MKC8866 in the same GB cells led to an attenuation of the expression of 490 of those proteins (**Fig. 1B**). Less that 10% of them were found to be modulated when the total proteome was analyzed. Interestingly, these proteins whose expression is IRE1-dependent exhibited steady mRNA expression levels (**Fig. 1C**). We next analyzed the expression of cells surface molecules in parental and RADH85/87 and U251 GB cells expressing a dominant negative form of IRE1 using a cell surface biotinylation approach (**Fig. 1D**). Again, the cell surface expression of 223 proteins was down-regulated after inhibition of IRE1 activity, most of those were associated with cell adhesion and migration (**Fig. 1E**), and exhibited an unaltered mRNA expression (**Fig. 1F**). These results highlight that IRE1 RNase activity promotes the secretion of multiple cell adhesion and cell-migration-related proteins.

**Figure 1.**
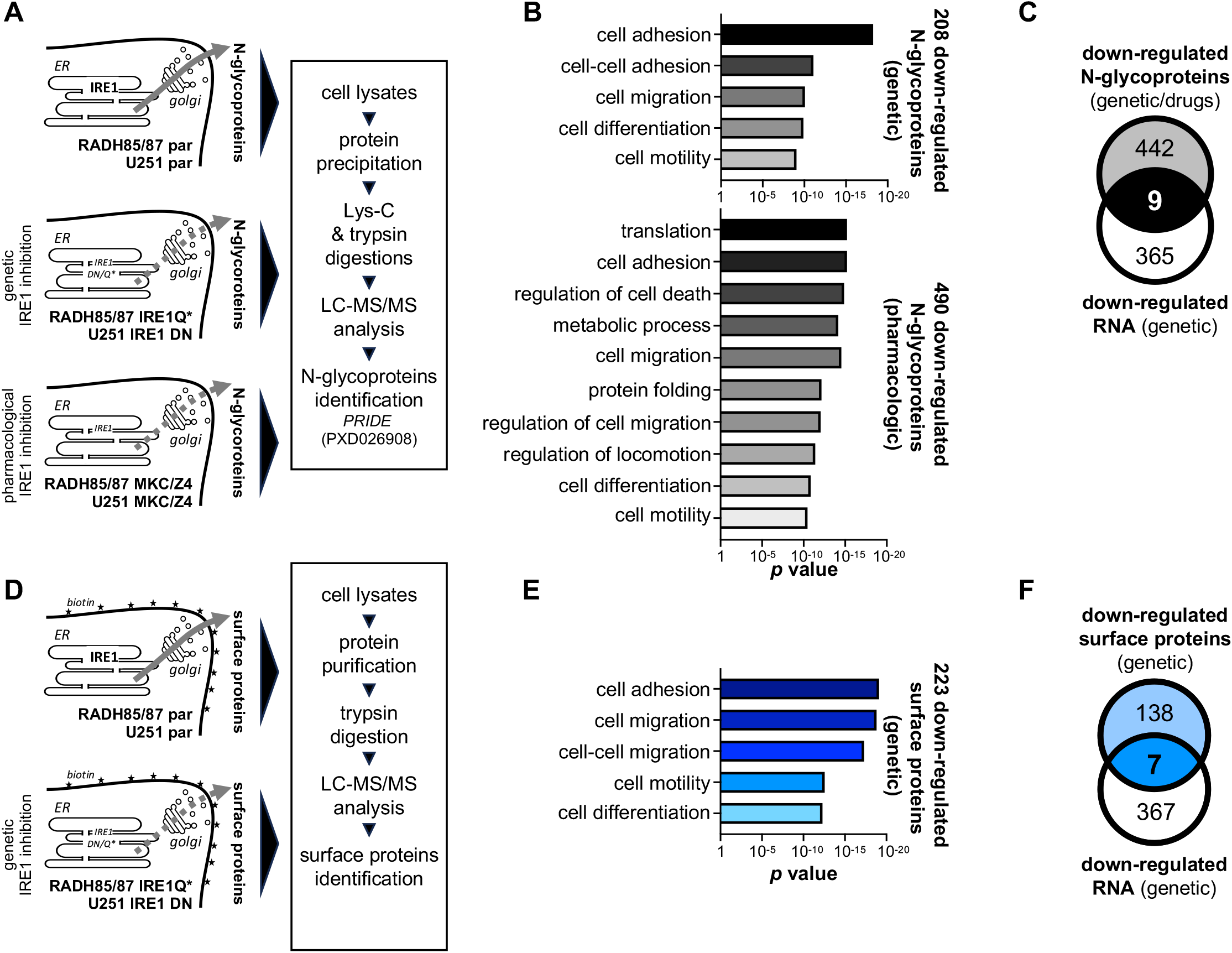
IRE1-dependent regulation of secreted proteins in GB cells. **(A)** A schematic representation of the proteomic approach used for analyzing N-glycoproteins present in IRE1-modulated GB cell lines U251, RADH85 and RADH87. IRE1 inhibition was performed genetically using overexpression of dominant negative IRE1 (IRE1_DN or IRE1_Q*) or pharmacologically using the IRE1 inhibitors MKC8866 or Z4. **(B)** Down-regulated N-glycoproteins were annotated using STRING resource (39). **(C)** Down-regulated N-glycoproteins were represented in a Venn diagram with genes which RNA expressions were down-regulated upon IRE1 inhibition. **(D)** A schematic representation of the proteomic approach used for analyzing cell surface proteins present in IRE1-modulated GB cell lines U251, RADH85 and RADH87. IRE1 inhibition was performed genetically using overexpression of dominant negative IRE1 (IRE1_DN or IRE1_Q*). **(E)** Down-regulated cell surface proteins were annotated using STRING resource. **(F)** Down-regulated cell surface proteins were represented in a Venn diagram with genes which RNA expressions were down-regulated upon IRE1 inhibition.

### IRE1 endoribonuclease activity promotes the expression of the Golgi associated protein GOLIM4 in GB cells

As IRE1 invalidation modulated the expression of secretory proteins without affecting their mRNA levels, we postulated that IRE1 could, instead of regulating the expression of the proteins, rather control important molecular actors involved in protein secretion and/or traffic. We therefore defined potential candidates as being functionally involved in protein secretion (and localizing to organelle of the SP) that were potential XBP1s targets, based on the ChIP-Atlas resource (29), and whose expression was down-regulated upon IRE1 inhibition (genetic or pharmacologic) in GB cells (**Fig. 2A**). Using these criteria, we identified 57 candidates (**Fig. 2A**). We selectively focused on 5 genes that were also potentially regulated by IRE1 in GB specimens based to our iRE1 activity signature described in (6), and whose expression modulation was associated with GB patients’ survival advantages or disadvantage (**Fig. 2B**). Messenger RNA expression of these 5 candidates was tested in 4 GB cell lines (RADH85/87, U251 and U87) in which the expression of IRE1 was attenuated using siRNA-mediated silencing (**Fig. S1A**). Amongst the tested candidates, only GOLIM4 mRNA expression was down-regulated in all GB cells treated with siRNA targeting IRE1 (**Fig. 2C**). We confirmed that GOLIM4 protein expression was reduced in GB cells upon IRE1 silencing (**Fig.S1B, 2D**) excepted in the RADH85 GB line (**Fig.S1C**). In addition, both IRE1 inhibitor MKC8866 (**Fig.S1D, 2D**) or XBP1 siRNA (**Fig.S1E, 2D**) led to reduced GOLIM4 protein expression in RADH87, U87 and U251 cells. GOLIM4 (for Golgi integral membrane protein 4, also named GPP130) is localized in the cis-Golgi, and is constitutively cycling between early endosomes, the Golgi and the plasma membrane. Overall, these data show that the expression of GOLIM4 protein is induced upon activation of the IRE1/XBP1s signaling axis in most of the GB cells tested in the present study.

**Figure 2.**
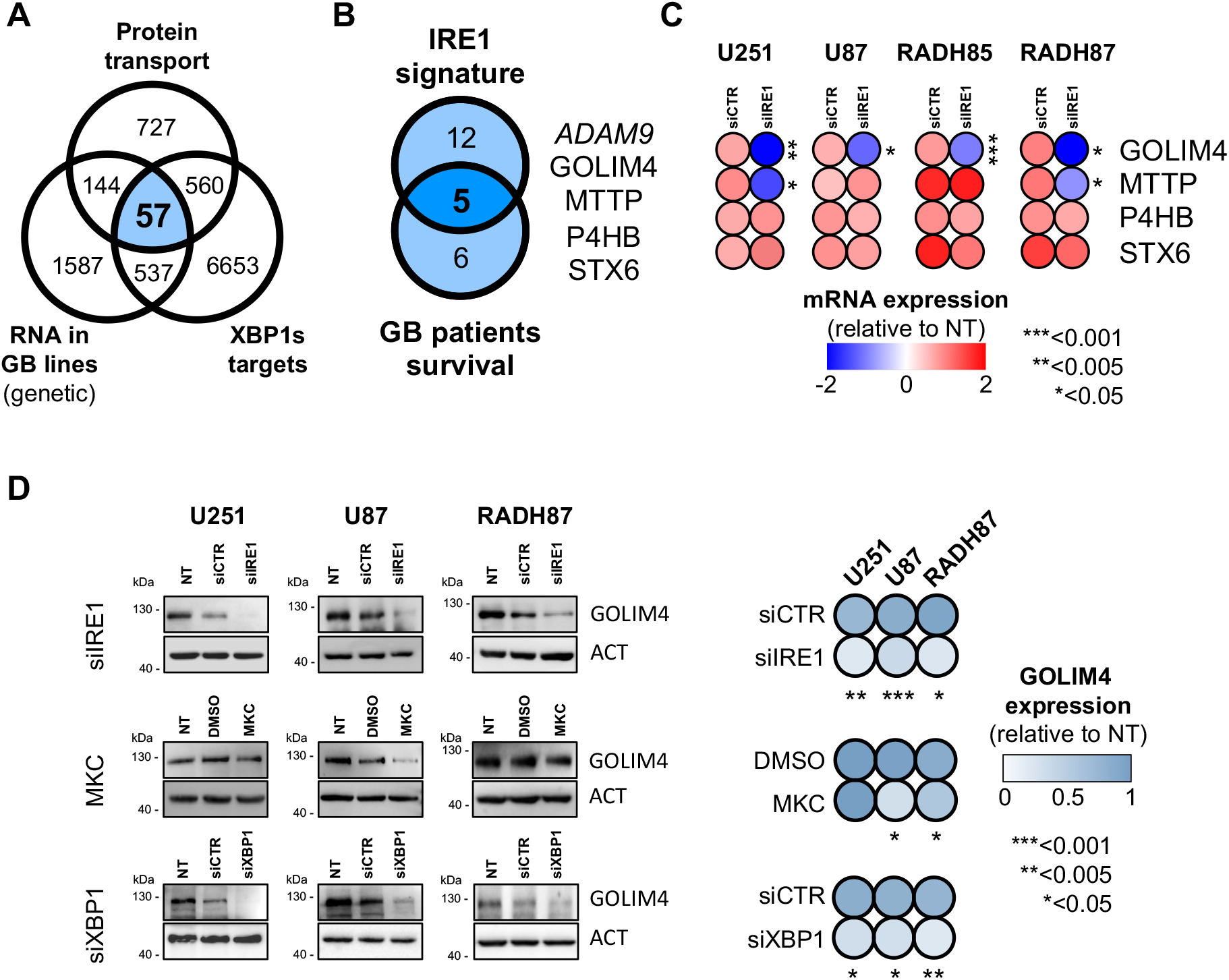
IRE1/XBP1s-dependent regulation of GOLIM4 in GB cells. **(A)** Genes involved in protein transport (from AMIGO gene ontology resource (40) GO:0015031) were represented in a Venn diagram with genes which RNA expressions were down-regulated upon IRE1 inhibition and putative genes regulated by XBP1s (from ChIP-Atlas resource (29)). **(B)** The selected genes were represented in a Venn diagram with genes modulated by IRE1 in GB specimens from the TCGA GB cohort (6) and GB patients survival. **(C)** mRNA expression of GOLIM4, MTTP, P4HB and STX6 from parental (NT), control (siCTR) and IRE1-silenced (siIRE1) GB cells U251, U87, RADH85 and RADH87 were analyzed by RT-Q-PCR. Expression levels were relative to parental cells (n=3); (*): *p*<0.05, (**): *p*<0.005 and (***): *p*<0.001. **(D)** Protein expression of GOLIM4 from parental (NT), control (siCTR or DMSO) and IRE1-modulated (siIRE1 or siXBP1) GB cells U251, U87 and RADH87 were analyzed by western-blot. IRE1 and XBP1s inhibition was performed using specific siRNA or the IRE1 inhibitor MKC8866 (MKC). Expression levels were relative to parental cells (n=3); (*): *p*<0.05, (**): *p*<0.005 and (***): *p*<0.001.

### IRE1 and GOLIM4 promote the secretion of N-glycosylated and cell surface proteins in GB cells

To evaluate the impact of GOLIM4 on protein secretion in our experimental systems, we first identified whose expression at the cell surface (cell surface biotinylation and mass spectrometry sequencing of the biotinylated proteins) was affected by IRE1 genetic or pharmacologic inhibition. This led to the identification of 25 cell surface proteins with reduced presence at the cell surface in at least 2 out 3 GB cell lines tested (**Fig. 3A**), whereas their mRNA expression remained unchanged. We then tested whether GOLIM4 silencing could also affect cell surface expression of those candidates. Importantly, siRNA-mediated GOLIM4 silencing (**Fig. 3B**) led also to reduce surface expression of MHC class I, IL13RA2, ITGB1 (a membrane partner of ITGA1/2), NCAM1 (CD56) and PDGFRA (CD140a) molecules, as determined by flow cytometry (**Fig. 3C**). Furthermore, others surface molecules involved in cell adhesion and migration including CD44, CD54 (ICAM1), CD90 and CD109 were also affected by the modulation of GOLIM4, at least in one of the two of the GB cell lines tested (**Fig.S2**).

**Figure 3.**
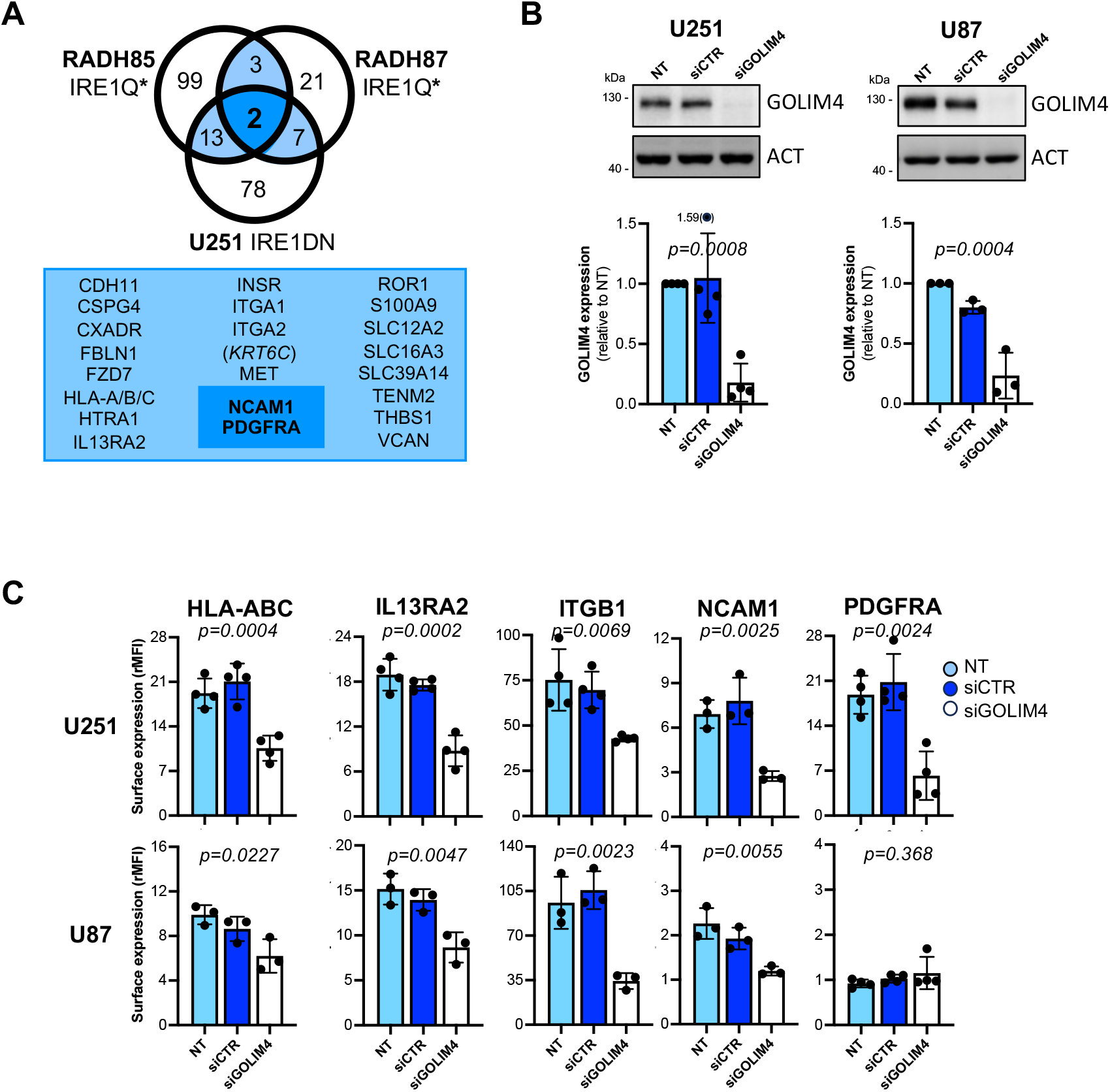
IRE1 and GOLIM4-dependent regulation of N-glycoproteins and surface proteins in GB cells. **(A)** Down-regulated surface proteins upon IRE1 genetic inhibition obtained from proteomic analyses of U251, RADH85 and RADH87 GB cells were represented in a Venn diagram. **(B)** GOLIM4 protein expression in parental (NT), control (siCTR) and GOLIM4-silenced (siGOLIM4) U251 and U87 GB cells was analyzed by western-blot. Protein expression levels were determined using Fiji; (n=3); *p* values were indicated on the top of the graph. **(C)** Surface expression of HLA-ABC, IL13RA2, ITGB1, NCAM1 and PDGFRA of parental (NT), control (siCTR) and GOLIM4-silenced (siGOLIM4) U251 and U87 GB cells was analyzed by flow cytometry. Protein expression levels were determined by the ratio of fluorescence intensity mean (n=3 to 4); *p* values were indicated on the top of the graph.

### GOLIM4 modulation affects cell-cell adhesion and migration of GB cells

As GOLIM4 controlled the presence of specific proteins at the cell surface and that those proteins were shown to be involved in cell adhesion and migration (without affecting their mRNA expression), we next analyzed the impact of the reduction of GOLIM4 on GB cell functions including cell proliferation, cell adhesion to extracellular matrix, cell-to-cell adhesion and cell migration.

GOLIM4 modulation with specific siRNA did not affect GB cell proliferation (**Fig.S3A**); cell adhesion to collagen, fibronectin and matrigel composed of multiple extracellular matrix proteins (**Fig.S3B**); or cell migration using a wound healing assay (**Fig.S3C**). However, down-regulation of GOLIM4 expression reduced cell-cell adhesion accessed by a cell aggregation assay (**Fig. 4A**) and increased GB cell migration in a Boyden chamber assay (**Fig. 4B**).

**Figure 4.**
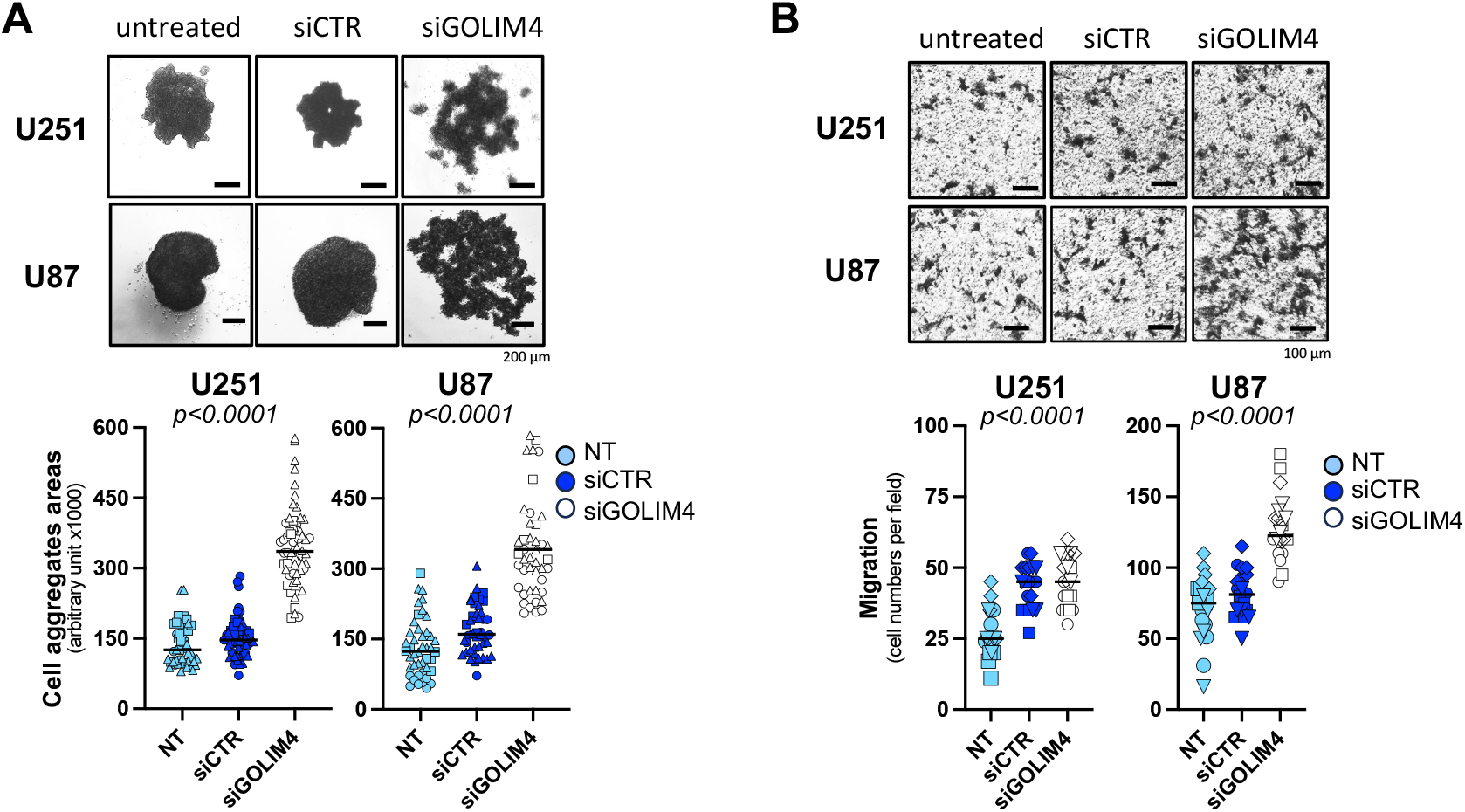
GOLIM4-dependent regulation of GB cell-to-cell adhesion and migration. **(A)** Parental (NT), control (siCTR) and GOLIM4-silenced (siGOLIM4) U251 and U87 GB cells was analyzed in a cell aggregation assay for testing cell-to-cell adhesion. Surface of cell aggregates were calculated using Fiji. High aggregate’s size corresponds to the low cell-to-cell adhesion (n=3); *p* values were indicated on the top of the graph. **(B)** Parental (NT), control (siCTR) and GOLIM4-silenced (siGOLIM4) U251 and U87 GB cells was using in a Boyden chamber migration assay. Migration was quantified by the mean of number of migrated cells observed per field (n=3; 5 fields per experiments); *p* values were indicated on the top of the graph.

Overall, these results demonstrate a novel IRE1/XBP1s/GOLIM4 regulon that controls cell surface and secreted proteins including adhesion molecules leading to affect cell-to-cell adhesion and migration in GB.

## Discussion

In the past years, we have demonstrated the important role of the ER stress sensor IRE1 in GB (18,19,23), in particular in the tumor microenvironment remodeling by affecting the angiogenesis (7) and immune infiltration (5), favoring the secretion of growth factors, cytokines and chemokines. In this context, we have hypothesized that IRE1 could more generally influence the protein secretion by directly impacting on the molecular machinery of the SP. In this study, we show that the activation of the IRE1/XBP1s branch of the UPR induces GOLIM4 expression, a Golgi-associated protein, that in turn modulates the presence of specific cell surface proteins involved in adhesion/migration of GB cells, hence identifying a new molecular link between IRE1 activation of cell adhesion/migration.

The molecule GOLIM4 (for Golgi integral membrane protein 4), also named GPP130 (for phosphor-protein 130 kDa) localizes to the cis-Golgi, and constitutively cycles between early endosomes, the Golgi and the plasma membrane (30,31). Lumenal pH of cellular compartments controls GOLIM4 distribution, since acidic pH causes GOLIM4 localization into early endosomes (30,32,33). GOLIM4 is involved in sorting both invasive toxins from *Shigella genus* (34) and endogenous secreted proteins into the early endosome-to-Golgi route (30). Importantly, manganese induces GOLIM4 oligomerization via its sortilin-mediated addressing into lysosomes for degradation (30,35,36). More recently, GOLIM4 has been described as a novel oncogene in human head and neck cancer by impacting on tumor cell division and resistance to apoptosis (37). GOLIM4 is also part of the chromosomal 3q amplicon found in multiple tumor types including head and neck, lung, and esophagus cancers; that encodes key regulators of secretory vesicles driving the tumor secretory addiction (38). In this work, we demonstrate that GOLIM4 down-regulation affects surface expression of molecules including those involved in cell adhesion and migration. One could speculate that GOLIM4 might contribute to the recycling of cell surface proteins by extracting them from the degradative route. This trafficking route from the early endosomes back into the Golgi apparatus might facilitate the maintenance of a certain protein level at the cell surface. Alternatively, GOLIM4 could directly act at the Golgi-to-secreted vesicles interface to mediate protein export to the plasma membrane. The precise mechanisms by which these targets are selected by GOLIM4 need to be further explored to better understand consequences on GB cell biology. Of note at the Golgi lumen, GOLIM4 bridges the calcium channel ATP2C1 to the Golgi phosphoprotein 3 GOLPH3 to allow the loading of calcium/CAB45-associated cargoes and the vesicle scission from the Golgi membrane (38). These molecular partners could participate to the selectivity of surface proteins recycled by GOLIM4.

In previous studies, the ER stress sensor IRE1 has been involved in regulating molecular actors of the SP associated with the ER and the ERGIC compartments. Through XBP1s induction, IRE1 induces COPII components SEC22, SEC23 and SEC24 (22,23), the ERGIC associated cargo receptor KDELR (23,24) and COPI molecules COPA, COPB1/2, COPE and COPG (23). This work adds new inputs on how IRE1 could affect protein secretion by modulating the protein trafficking between the plasma membrane, the early endosomes and the Golgi apparatus. Furthermore, this novel IRE1-dependent level of protein secretion of surface molecules such as ITGB1 and NCAM1 and directly impacts important GB functions including cell adhesion and migration. The current work implies that targeting IRE1 signaling might impede GB aggressiveness by attenuating tumor secretory functions that influence tumor cell invasion, inflammation and immunity (5).

## Author Contributions

KB, MN, SM, LN – methodology, investigation, formal analysis; EL, EC – conceptualization, funding acquisition, writing (review and editing); TA – supervision, conceptualization, methodology, investigation, formal analysis, funding acquisition, project administration, writing (original draft, review and editing) (https://www.casrai.org/credit.html).

