## Supplemental_Materials for "IRE1-dependent GOLIM4 expression controls protein secretion to modulate glioblastoma cell adhesion and migration"

#### TABLE OF CONTENTS


SUPPLEMENTAL METHODS AND MATERIALS..…...………………..…… page 2
TABLE S1 ………………………………………………………………… page 3
TABLE S2 ………………………………………………………...…………. page 4
SUPPLEMENTAL FIGURE LEGENDS ………………………………….. page 5
FIGURE S1 ……………………………………………………...………… page 6
FIGURE S2 ……………………………………………………...………… page 7
FIGURE S3 ……………………………………………………...………… page 8

#### SUPPLEMENTAL METHODS AND MATERIALS

***Cell proliferation –*** Parental, control and GB cells silenced for GOLIM4 were cultured in 96-well plates at the concentration of $1x10^3$ cells/well and the growth rate was measured daily for 6 days using the Incucyte apparatus (SX5 Live-Cell Analysis System, Essen BioScience, Germany). Phase images were taken every 24 hours and were recorded (400 ms exposure, 10x lens) for each condition in triplicate. Images were analyzed using the Adherent Cell-by-cell on phase algorithm from Incucyte 2022B Rev2 software. The proliferation index was given by the ratio of the specific index observed with tested cells and the specific index observed with cells used at the beginning of the experiment.

***Cell adhesion to extracellular matrix –*** Ninety-six-well plates were coated with a filtered solution of 400 µg/mL rat tail collagen I, 1 mg/mL fibronectin and 1% matrigel solution (BD Biosciences) in PBS. Parental, control and GOLIM4-silenced U251 and U87 cells (25,000 cells) were plated for time points 0, 15, 30 60 and 120 minutes. Medium and unattached cells were aspirated. Wells were washed with PBS and attached cells were stained with the WST1 reagent. The percentage of cell attachment was calculated by the ratio of the specific OD observed with tested cells and the specific OD observed with total cells used at the beginning of the experiment.

***Wound healing migration assay –*** Parental, control and GB cells silenced for GOLIM4 were cultured in 96-well plates at the concentration of 30,000 cells/well for 24 hours. Wound on the cell monolayers was applied using a wound maker (Essen BioScience). The cell migration rate was measured daily for 48 hours using the Incucyte apparatus. Phase images were taken every 2 hours and were recorded (400 ms exposure, 10x lens) for each condition in triplicate. Images were analyzed the Schrach Wound algorithm from Incucyte 2022B Rev2 software.

#### SUPPLEMENTAL TABLES

**Table S1. Antibodies used in the study**

| Targets | Species / isotype | Fluo. | Clone / RRID | Compagny |
|---|---|---|---|---|
| *for western-blot* | | | | |
| **ACTIN** | mouse mAb | | AC-74 / AB_476743 | Sigma Aldrich |
| **GOLIM4** | mouse mAb | | XY-2 / AB_2247835 | Santa Cruz |
| **IRE1** | rabbit mAb | | 14C10 / AB_823545 | Cell Signaling |
| **Targets** | **Species / isotype** | **Fluo.** | **Clone / RRID** | **Compagny** |
| *for flow cytometry* | | | | |
| **controls** | mouse mAb (IgG1) | FITC | MOPC21 / AB_2891079 | BioLegend |
| | mouse mAb (IgG2a) | FITC | MOPC173 / AB_2884007 | BioLegend |
| | mouse mAb (IgG2b) | FITC | 27-35 / AB_396085 | BD Biosciences |
| | mouse mAb (IgG1) | PE | X40 / AB_400130 | BD Biosciences |
| | mouse mAb (IgG2a) | PE | G155-178 / AB_11151914 | BD Biosciences |
| | mouse mAb (IgG1) | APC | MOPC21 / AB_2888687 | BioLegend |
| | mouse mAb (IgG2b) | APC | 27-35 / AB_398612 | BD Biosciences |
| | mouse mAb (IgG1) | BV421 | A85-1 / AB_2737664 | BD Biosciences |
| **CD44** | mouse mAb (IgG2b) | APC | G44-26 / AB_398683 | BD Biosciences |
| **CD109** | mouse mAb (IgG1) | PE | TEA 2/16 / AB_396311 | BD Biosciences |
| **HLA-ABC** | mouse mAb (IgG2a) | FITC | W6/32 / AB_2566253 | BioLegend |
| **ICAM1 (CD54)** | mouse mAb (IgG1) | APC | HA58 / AB_395901 | BD Biosciences |
| **IL13Rα2 (CD213a2)** | mouse mAb (IgG1) | APC | SHM38 / AB_2562583 | BioLegend |
| **ITGB1 (CD29)** | mouse mAb (IgG1) | BV421 | MAR4 / AB_2741751 | BD Biosciences |
| **NCAM1 (CD56)** | mouse mAb (IgG2b) | FITC | NCAM16.2 / AB_397180 | BD Biosciences |
| **PDGFRA (CD140a)** | mouse mAb (IgG2a) | PE | αR1 / AB_2737804 | BD Biosciences |
| **THY1 (CD90)** | mouse mAb (IgG1) | FITC | 5E10 / AB_893429 | BioLegend |

**Table S2. Primers used in the study**

| Gene | Forward primer | Reverse primer |
|---|---|---|
| **ACTIN** | 5'-CATGGGTGGAATCATAATGG-3' | 5-AGCACTGTGTTGCGCTACAG-3' |
| **GAPDH** | 5'-AAGGTGAAGGTCGGAGTCAA-3' | 5'-CATGGGTGGAATCATAATGG-3' |
| **GOLIM4** | 5'-ATGAGCCTCGTGAACAAGGACC-3' | 5'-CTCTCACTTGCTCGGCTTCTTC-3' |
| **IRE1** | 5'-GCCACCCTGCAAGAGTATGT-3' | 5'-ATGTTGAGGGAGTGGAGGTG-3' |
| **MTTP** | 5'-AGGCTGTCAGAAACTTCCTGGC-3' | 5'-GTCTGAGCAGAGGTGACAGCAT-3' |
| **P4HB** | 5'-AGGCTGATGACATCGTGAACT-3' | 5'-GGTATTTGGAGAACACGTCACTG-3' |
| **STX6** | 5'-CACGAATTGGAGAGCACTCAGTC-3' | 5'-GAGGATGAGCACAACCAACAGG-3' |
| **XBP1s** | 5'-TGCTGAGTCCGCAGCAGGTG-3' | 5'-GCTGGCAGGCTCTGGGGAAG-3' |

#### SUPPLEMENTAL FIGURE LEGENDS

**Figure S1. IRE1/XBP1s-dependent regulation of GOLIM4 in GB cells**

**(A)** Parental (NT), control (siCTR) and IRE1 silenced (siIRE1) U251, U87, RADH85 and RADH87 GB cells were tested for IRE1 mRNA down-regulation by RT-Q-PCR. mRNA expression levels were relative to parental cells (n=3 to 4); *p* values were indicated on the top of the graph. **(B)** Parental (NT), control (siCTR) and IRE1 silenced (siIRE1) U251, U87 and RADH87 GB cells were tested for IRE1 protein down-regulation by western-blot. Protein expression levels were relative to parental cells (n=3); *p* values were indicated on the top of the graph. **(C)** Parental (NT), control (siCTR) and IRE1 or GOLIM4 silenced (siIRE1 or siGOLIM4 respectively) RADH85 GB cells were tested for IRE1 and GOLIM4 protein down-regulation by western-blot. Protein expression levels were relative to parental cells (n=3); *p* values were indicated on the top of the graph. **(D and E)** Parental (NT), control (DMSO and siCTR) IRE1-modulated (MKC and siXBP1) U251, U87 and RADH87 GB cells were tested for XBP1s mRNA down-regulation by RT-Q-PCR. IRE1 inhibition was performed using MKC8866 (MKC) **(D)** or XBP1s silencing using specific siRNA **(E)**. mRNA expression levels were relative to parental cells (n=3 to 4); *p* values were indicated on the top of the graph.

**Figure S2. GOLIM4-dependent regulation of surface proteins in GB cells**

Surface expression of CD44, CD54 (ICAM1), CD90 (THY1) and CD109 of parental (NT), control (siCTR) and GOLIM4-silenced (siGOLIM4) U251 and U87 GB cells was analyzed flow cytometry. Protein expression levels were determined by the ratio of fluorescence mean (n=3 to 4); *p* values were indicated on the top of the graph.

**Figure S3. GOLIM4-dependent regulation of GB cell proliferation, cell adhesion to extracellular matrix and cell migration**

**(A)** Parental (NT), control (siCTR) and GOLIM4 silenced (siGOLIM4) U251 and U87 GB cells were tested for cell proliferation using the Incucyte apparatus. The proliferation index is given by the ratio of the specific index observed with tested cells and the specific index observed with cells used at the beginning of the experiment (n=3); *p* values were indicated on the top of the graph. **(B)** Parental (NT), control (siCTR) and GOLIM4 silenced (siGOLIM4) U251 and U87 GB cells were used in a cell adhesion to extracellular matrix assay with collagen (COL), fibronectin (FN) and matrigel substrates. Results were expressed as percentage of adherent cells (n=3); *p* values were indicated on the top of the graph. **(B)** Parental (NT), control (siCTR) and GOLIM4 silenced (siGOLIM4) U251 and U87 GB cells were tested in a wound healing assay for cell migration using the Incucyte apparatus. Results were expressed as percentage of migrating cells obtained with the Incucyte software (n=3); *p* values were indicated on the top of the graph.

#### FIGURE S1

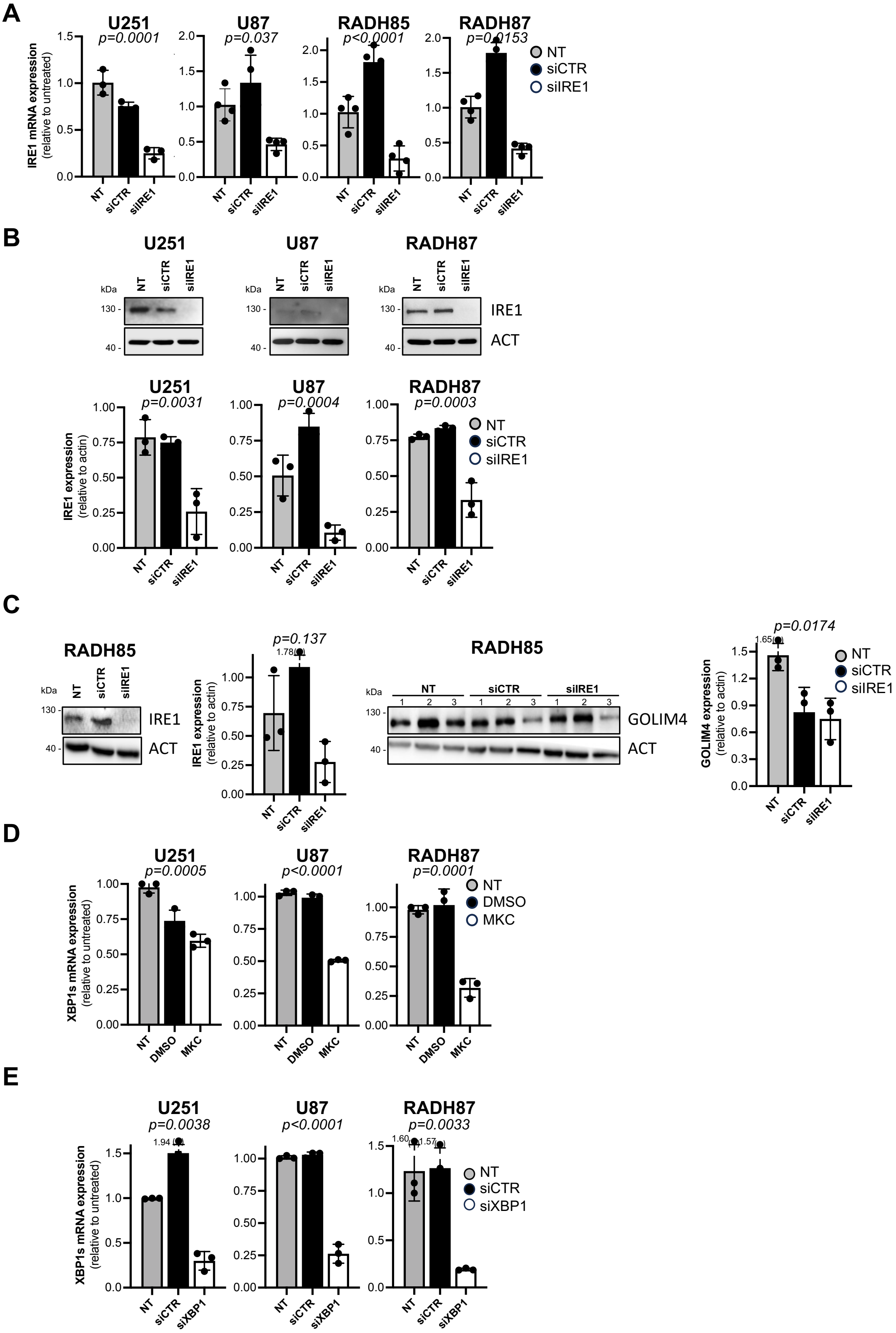
A
U251
p=0.0001
U87
p=0.037
RADH85
p<0.0001
RADH87
p=0.0153
IRE1 mRNA expression
(relative to untreated)
NT
siCTR
siIRE1
B
U251
U87
RADH87
kDa
130 -
40 -
IRE1
ACT
U251
p=0.0031
U87
p=0.0004
RADH87
p=0.0003
IRE1 expression
(relative to actin)
NT
siCTR
siIRE1
C
RADH85
IRE1
ACT
p=0.137
1.78
IRE1 expression
(relative to actin)
RADH85
NT
siCTR
siIRE1
GOLIM4
ACT
p=0.0174
1.65
GOLIM4 expression
(relative to actin)
D
U251
p=0.0005
U87
p<0.0001
RADH87
p=0.0001
XBP1s mRNA expression
(relative to untreated)
NT
DMSO
MKC
E
U251
p=0.0038
1.94
U87
p<0.0001
RADH87
p=0.0033
1.60
1.57
XBP1s mRNA expression
(relative to untreated)
NT
siCTR
siXBP1

### FIGURE S2

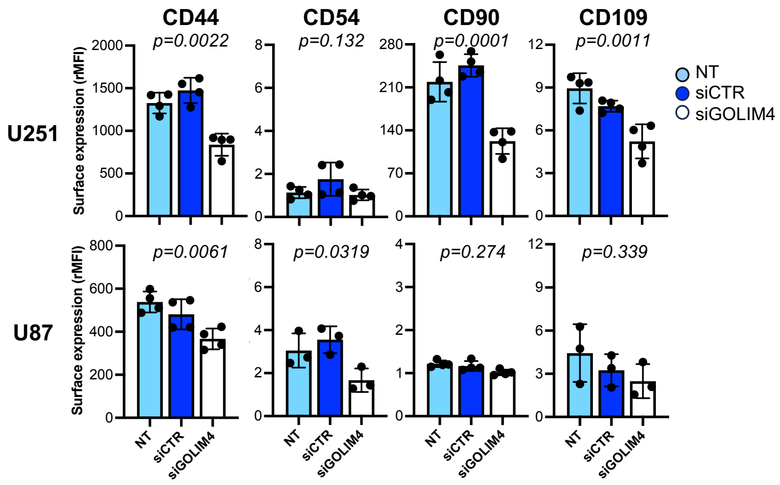
CD44
CD54
CD90
CD109
p=0.0022
p=0.132
p=0.0001
p=0.0011
p=0.0061
p=0.0319
p=0.274
p=0.339
U251
U87
Surface expression (rMFI)
NT
siCTR
siGOLIM4

### FIGURE S3

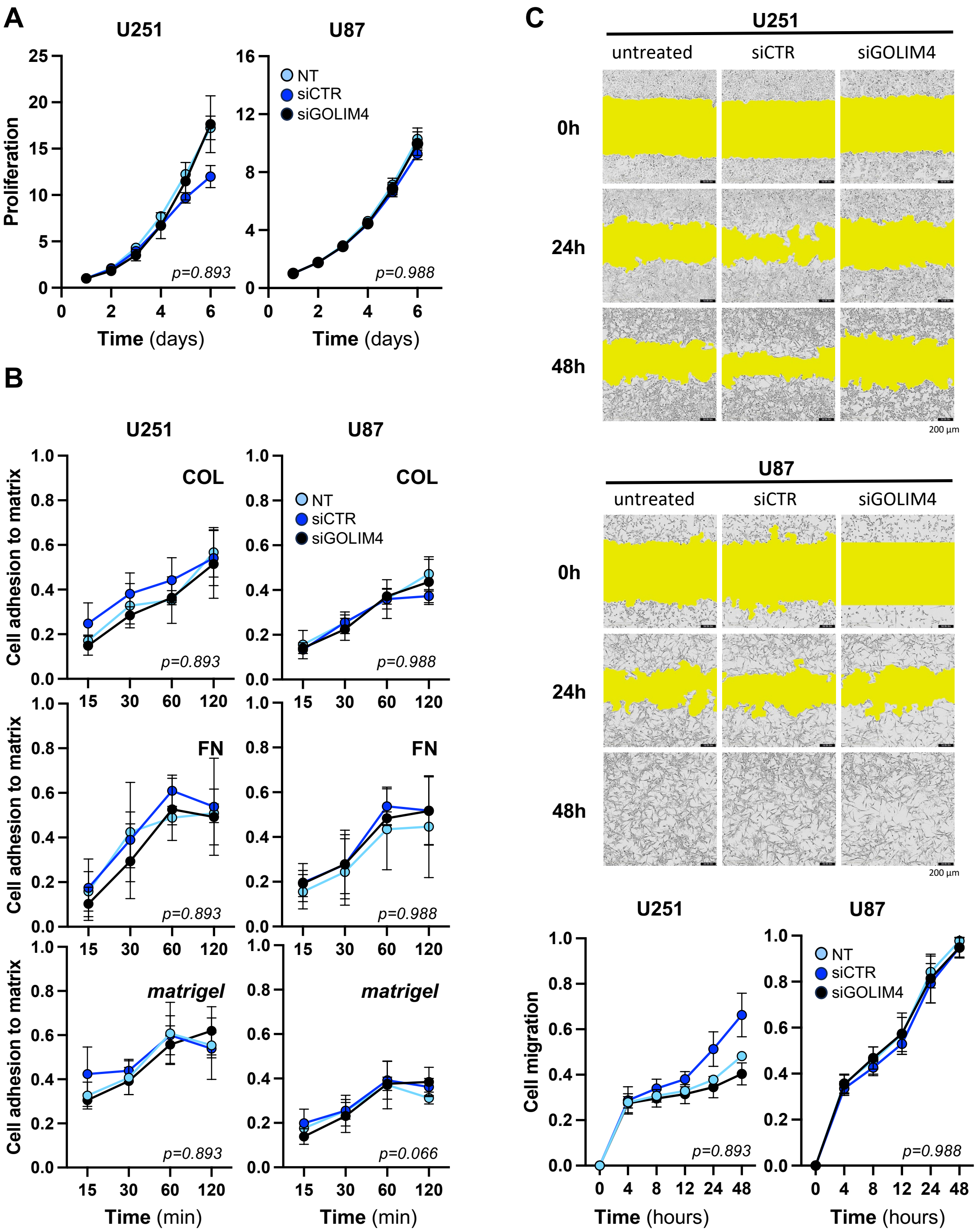
A
U251
U87
Proliferation
Time (days)
NT
siCTR
siGOLIM4
p=0.893
p=0.988
B
COL
FN
matrigel
Cell adhesion to matrix
Time (min)
p=0.066
C
untreated
siCTR
siGOLIM4
0h
24h
48h
200 µm
Cell migration
Time (hours)
